# Opposing functions of the Hda1 complex and histone H2B mono-ubiquitylation in regulating cryptic transcription initiation in *Saccharomyces cerevisiae*

**DOI:** 10.1101/2020.10.12.336081

**Authors:** Margaret K. Shirra, Rachel A. Kocik, Mitchell A. Ellison, Karen M. Arndt

**Author notes:** Corresponding Author: Karen M. Arndt, Department of Biological Sciences University of Pittsburgh, 4249 Fifth Avenue, Pittsburgh, PA 15260.

## Abstract

Maintenance of chromatin structure under the disruptive force of transcription requires cooperation among numerous chromatin regulatory factors. Histone post-translational modifications can regulate nucleosome stability and influence the disassembly and reassembly of nucleosomes during transcription elongation. The Paf1 transcription elongation complex, Paf1C, is required for several transcription-coupled histone modifications, including the mono-ubiquitylation of H2B. In *Saccharomyces cerevisiae*, amino acid substitutions in the Rtf1 subunit of Paf1C greatly diminish H2B ubiquitylation and cause transcription to initiate at a cryptic promoter within a coding gene, an indicator of chromatin disruption. In a genetic screen to identify factors that functionally interact with Paf1C, we identified a mutation in *HDA3*, a gene encoding a subunit of the Hda1C histone deacetylase, as a suppressor of an *rtf1* mutation. Absence of Hda1C also suppresses the cryptic initiation phenotype of other mutants defective in H2B ubiquitylation. The genetic interactions between Hda1C and the H2B ubiquitylation pathway appear specific: loss of Hda1C does not suppress the cryptic initiation phenotypes of other chromatin mutants and absence of other histone deacetylases does not suppress the absence of H2B ubiquitylation. Providing further support for an appropriate balance of histone acetylation in regulating cryptic initiation, we find that deletion of the Sas3 histone acetyltransferase elevates cryptic initiation in *rtf1* mutants. Our data suggest a coordination between two epigenetic modifiers, the H2B ubiquitylation pathway and Hda1C, in regulating chromatin structure during transcription elongation and reveal an unexpected role for a histone deacetylase in supporting chromatin accessibility.

## INTRODUCTION

Transcription of eukaryotic genes requires passage of RNA polymerase through a generally repressive chromatin template. Nucleosomes, which consist of approximately 147 base pairs of DNA wrapped around an octamer of histones H2A, H2B, H3 and H4, present a barrier to RNA polymerase II (Pol II) progression (Kujirai *et al*. 2018; Farnung *et al*. 2018; Chen *et al*. 2019). Organisms have evolved numerous mechanisms to overcome this barrier. Variants of histones can replace their canonical counterparts at certain locations in the genome, such as the presence of H2A.Z in exchange for H2A in the nucleosome downstream of the transcription start site (Bagchi *et al*. 2020). Chromatin remodeling factors, such as those in the SWI/SNF, ISWI, INO80, and CHD families, alter the positioning of nucleosomes to regulate DNA accessibility (Tyagi *et al*. 2016), and the histone chaperones FACT and Spt6 facilitate the reassembly of nucleosomes following Pol II passage (Gurard-Levin *et al*. 2014; Hammond *et al*. 2017). The post-translational modification of histones, though, may represent the most diversified means by which nucleosomes can regulate Pol II transcription.

With high selectivity, histone modifying enzymes carry out the covalent modification (*e.g.* phosphorylation, acetylation, methylation, or ubiquitylation) of amino acids within the N- and C-terminal tails or core domains of the histones (Bannister and Kouzarides 2011; Lawrence *et al*. 2016). At the genomic level, these modifications are deposited in specific patterns and, depending on their functions, are enriched in transcribed or non-transcribed regions. Enzymes that modify histones are hypothesized to generate a “histone code” that is interpreted by effector proteins, which bind with high specificity to appropriately modified histones (Jenuwein and Allis 2001). For example, acetylation of lysines by histone acetyltransferases (HATs) (Lee and Workman 2007) loosens the interactions of histone tails with the DNA backbone (Hong *et al*. 1993), providing increased DNA accessibility to the transcription machinery and modulating transcription. In addition, acetylated histones can serve as binding sites for other regulatory factors, such as chromatin remodeling factors (Musselman *et al*. 2012; Marmorstein and Zhou 2014). Histone deacetylases (HDACs) remove the acetyl groups, thus reversing these effects (Seto and Yoshida 2014; Porter and Christianson 2019; Park and Kim 2020).

Crosstalk among histone modifications allows the integration of multiple signals to control transcription. A modification may be a pre-requisite for another modification. One well-studied example is the dependency of H3K4 and H3K79 di- and tri-methylation (me2 and me3) on the mono-ubiquitylation of a conserved lysine in H2B (K123 in *S. cerevisiae*; K120 in *H. sapiens*) (Worden and Wolberger 2019). H2B K123 mono-ubiquitylation (H2Bub) is catalyzed by the ubiquitin conjugating enzyme Rad6 and the ubiquitin protein-ligase Bre1 (Robzyk *et al*. 2000; Hwang *et al*. 2003; Wood *et al*. 2003a; Fuchs and Oren 2014) and is enriched on the bodies of actively transcribed genes (Minsky *et al*. 2008; Batta *et al*. 2011). The Set1 and Dot1 histone methyltransferases that modify H3K4 and H3K79, respectively, require the ubiquitin moiety on H2B for activity (Briggs *et al*. 2002; Ng *et al*. 2002; Sun and Allis 2002; Dover *et al*. 2002; Hsu *et al*. 2019; Worden and Wolberger 2019; Worden *et al*. 2020). In turn, H3K4 methylation as well as H3K36me3 recruit the NuA3 HAT to the genome through its Yng1 and Pdp3 subunits, thereby controlling the location of histone acetylation by the Sas3 subunit

(Taverna *et al*. 2006; Martin *et al*. 2017). In other cases, one histone modification may regulate the removal of another modification. For example, H3K36me2/3 by Set2 activates Rpd3(S), a deacetylase complex, to remove acetyl groups from histones and restore a repressive chromatin structure that prevents initiation of Pol II transcription within protein coding regions (Carrozza *et al*. 2005; Joshi and Struhl 2005; Keogh *et al*. 2005; Drouin *et al*. 2010; Govind *et al*. 2010; Ruan *et al*. 2015; Venkatesh and Workman 2015). Thus, the enzymes that modify histones coordinately control chromatin structure and collectively impose or remove barriers to transcription.

In addition to the histone modifiers themselves, transcription factors that associate with Pol II can significantly influence the epigenetic state of chromatin. Previous work on the *S. cerevisiae* Polymerase-Associated Factor 1 Complex (Paf1C), which is composed of Paf1, Ctr9, Cdc73, Rtf1, and Leo1, has demonstrated a key role for this transcription elongation complex in regulating conserved transcription-coupled histone modifications. Paf1C has not been shown to possess enzymatic activity; however, mutations in genes encoding Paf1C subunits greatly diminish the levels of H2Bub, H3K4me2/3, H3K79me2/3 and H3K36me3 (Krogan *et al*. 2003; Ng *et al*. 2003a, 2003b; Wood *et al*. 2003b; Laribee *et al*. 2005; Chu *et al*. 2007; Piro *et al*. 2012; Van Oss *et al*. 2016, 2017). We previously identified a small region within the Rtf1 subunit of Paf1C, termed the histone modification domain (HMD), that appears to function as a critical cofactor in the deposition of H2Bub. The HMD is necessary and sufficient to promote H2Bub *in vivo*, stimulates H2Bub *in vitro*, and binds directly to Rad6 (Piro *et al*. 2012; Van Oss *et al*. 2016). Mutations that disrupt the Rad6-HMD interaction severely reduce levels of H2Bub, H3K4me2/3 and H3K79me2/3 and cause phenotypes indicative of disrupted chromatin structure (Warner *et al*. 2007; Tomson *et al*. 2011; Van Oss *et al*. 2016). For example, certain amino acid substitutions within the HMD lead to transcription initiation at a cryptic transcription start site (TSS) within the coding region of a Pol II transcribed gene (Tomson *et al*. 2011; Van Oss *et al*. 2016), a mutant phenotype that arises when the repressive chromatin state of transcribed regions is disrupted by the loss of histone chaperones, histone modifiers, and chromatin remodeling factors (Kaplan *et al*. 2003; Cheung *et al*. 2008; Silva *et al*. 2012). In this study, we exploited the cryptic transcription initiation phenotype of HMD mutants and performed a genetic screen to investigate the functional interactions involving this domain and the histone modification it facilitates, H2Bub. Our results uncovered an interplay between the factors that establish H2Bub during transcription elongation and the Hda1 deacetylase complex (Hda1C) and argue that Hda1C impacts chromatin accessibility within gene bodies.

## MATERIALS AND METHODS

### Yeast strains and growth conditions

*Saccharomyces cerevisiae* strains were grown at 30 °C in YPD or SC media (Rose *et al*. 1990). The SC-His+Gal medium used to monitor the cryptic initiation phenotype contained 2% galactose. To assess defects in telomeric silencing, SC media contained 0.1% 5-fluoroorotic acid (5-FOA; USBiological, F5050). *S. cerevisiae* strains used in this study are listed in Table S1. KY strains are isogenic with FY2, a *GAL2^+^* derivative of S288C (Winston *et al*. 1995). KA strains were derived from crosses originating with strains from a library of histone H3 mutants (Dai *et al*. 2008). Strains were made by standard methods for genetic crosses or gene replacements (Rose *et al*. 1990; Vidal *et al*. 1991). Strains containing a mutation in *RTF1* were identified during tetrad analysis by the presence of an HA-tag sequence, using PCR amplification. Strains containing the *htb1-K123R* allele were identified by restriction enzyme digestion as previously described (Tomson *et al*. 2011). For phenotypes assessed by replica plating, strains were first purified on YPD. For serial dilution analysis, strains were grown overnight in liquid YPD, diluted to OD_600_ = 0.8, and 3 µl of five-fold serial dilutions were spotted onto SC-His+Gal or SC Complete media.

### Identification of a mutation that suppresses defects in the Rtf1 HMD

With the original goal of identifying high-copy-number suppressors of the *rtf1-108-110A* mutation, KY1232 was transformed with a *LEU2*-marked high-copy-number plasmid library and replica-plated to SC-His-Leu+Gal and SC-Leu+5FOA media to monitor suppression of the cryptic initiation and telomeric silencing phenotypes, respectively.

Using a strategy previously described (Thompson *et al*. 1993), the pRS425-based (Christianson *et al*. 1992) library was made in our laboratory from genomic DNA prepared from an *rtf1Δ* strain (KY957) to avoid recovering *RTF1* plasmids that complemented the *rtf1-108-110A* mutation. Two 5FOA resistant (5FOA^R^), His^-^ strains were isolated and, surprisingly, retained the suppression phenotypes even after plasmid loss through growth in nonselective conditions, indicating that they harbored chromosomal suppressors of *rtf1-108-110A*. The two strains, following plasmid loss, were saved as KY1410 and KY1411 and studied further. Diploid yeast from genetic matings with *rtf1-108-110A* strains and KY1410 and KY1411 showed that the two suppressor mutations were recessive, as the diploids grew on SC-His+Gal media. Additional crosses between KY1410 and KY1411 and with a strain containing *RTF1*, KY1228, showed that the suppression phenotype of each strain was due to a mutation in a single gene and that the suppressor mutation in each strain was unlinked to the *rtf1-108-110A* mutation. Haploid derivatives from these crosses were used in genetic matings to show that the two suppressor mutations defined a single complementation group. Strains containing the *sup2-22* mutation were chosen for further analysis.

Several attempts to identify the suppressor mutation by complementation of the cryptic initiation phenotype using plasmid-based yeast genomic DNA libraries were unsuccessful. Two genes that reversed the cryptic initiation phenotype of the *rtf1-108-110A sup2-22* strain were *HSF1* and *ASF1*. However, further genetic analysis showed that the suppressor mutation was not in these genes, but rather expression of these genes from a plasmid was likely affecting expression from the cryptic initiation reporter. Specifically, in the *FLO8* gene, upstream of where the *HIS3* reporter is inserted, we noticed a potential Hsf1 binding site 80 bp upstream of the predicted TATA box for the cryptic transcript, suggesting that overexpression of Hsf1 may drive higher expression of the *HIS3* reporter. Further, *RTF1* strains that were transformed with an Asf1-expressing plasmid grew on the SC-His+Gal media, suggesting that Asf1 was bypassing the effect of the suppressor. This is consistent with the role of Asf1 in promoting histone exchange and increasing histone acetylation on gene bodies (Hammond *et al*. 2017; Zhang *et al*. 2018). Therefore, we performed bulk segregant analysis (Birkeland *et al*. 2010) followed by whole genome sequencing to identify the *sup2-22* suppressor.

Six tetrads were selected from a cross between KY1232 and KY1498, and genomic DNA was prepared (Gopalakrishnan and Winston 2019) from the two pools of twelve haploid progeny either exhibiting a *sup2-22* phenotype or not exhibiting a *sup2-22* phenotype. Sequences were obtained from libraries prepared using an NEB Ultra II Library Prep Kit (New England Biolabs # E7103), multiplexed and run on an Illumina MiSeq with a 150-cycle v3 Reagent Kit yielding approximately 7.6 million reads per sample. Raw reads were mapped to the *S. cerevisiae* genome (S288C version = R64-2-1) (Cherry *et al*. 2012; Engel *et al*. 2014) using Bowtie2 (Langmead and Salzberg 2012). SAMtools was used to calculate genotype likelihoods before calling sequence variants using BCFtools (Li *et al*. 2009; Li 2011). Using VCFtools (Danecek *et al*. 2011), VCF files for mutant and wild-type segregants were compared and only the mutations found in the suppressor pool were selected from the output text (Ellison *et al*. 2020).

Variants with a quality score > 50 and found within coding regions were retained, leaving 3 variants. Two of the variants were synonymous (Mtl1 S257S and Flo5 V354V), and a third variant, which created a frameshift (GT to C at +1118), mapped within the *HDA3* gene. Sequence variants of interest were further inspected by visualizing the data in the Integrative Genomics Viewer (IGV) from the Broad Institute (Thorvaldsdottir *et al*. 2013). The mutation at codon 373 of *HDA3* is predicted to change seven amino acids in the protein product before a stop codon is encountered.

### Western blot analysis

Western blots to visualize H2Bub were performed using whole-cell extracts prepared using SUTEB buffer as described (Van Oss *et al*. 2016). H3 methylation was examined using whole-cell extracts prepared using a TCA extraction method (Van Oss *et al*. 2016). Proteins were run on 15% SDS-polyacrylamide gels. The following antibodies were used: α-H2B (Active Motif #39237; 1:3000), α-H2Bub (Cell Signaling #5546; 1:1000), α-G6PDH (Sigma #A9521; 1:20000), α-H3K4Me2 (Millipore #07-030; 1:2000), α-H3K4Me3 (Active Motif #39159; 1:2000), α-H3K79Me2/3 (Abcam #ab2621; 1:1000; note that this antibody recognizes both di- and trimethylated H3K79), and α-H3 antibody (Tomson *et al*. 2011). Blots were developed using Thermo Scientific West Pico Plus (34580) and images were collected on a BioRad ChemiDoc™ XRS+.

### Reagent and data availability

All strains and detailed protocols are available upon request. Sequencing data are available at NCBI as part of the BioProject: PRJNA634539, BioSample accessions: SAMN14999039, SAMN14999040.

## RESULTS

### Loss of Hda1C suppresses three different transcription-associated phenotypes of *rtf1* HMD mutants

In *S. cerevisiae*, the Rtf1 HMD is required for H2Bub through its direct interaction with Rad6 (Van Oss *et al*. 2016). To identify factors that functionally interact with the HMD, we conducted a genetic screen for suppressors of a previously characterized mutation, *rtf1-108-110A*, that replaces three consecutive residues within the HMD with alanine and greatly decreases global H2Bub levels *in vivo* (Tomson *et al*. 2011). Strains containing the *rtf1-108-110A* mutation exhibit transcription initiation from a cryptic TSS within the *FLO8* open reading frame (Tomson *et al*. 2011), as assayed by a *GAL1p-FLO8-HIS3* reporter construct (Cheung *et al*. 2008). *RTF1* strains carrying this reporter are His^-^ because transcription initiation at the canonical *FLO8* TSS gives rise to an mRNA in which the *HIS3* coding sequence is out of frame with respect to *FLO8*.

However, in mutants defective in maintaining proper chromatin architecture, such as the *rtf1-108-110A* mutant, transcription initiation occurs at a cryptic TSS within the *FLO8* gene, creating a transcript from which *HIS3* is expressed (Figure 1A). We exploited this phenotype to isolate genetic suppressors of *rtf1-108-110A* with the initial intent of identifying genes that, when overexpressed from a plasmid, would suppress the His^+^ phenotype of *rtf1-108-110A* mutants containing the *GAL1p-FLO8-HIS3* reporter.

**Figure 1:**
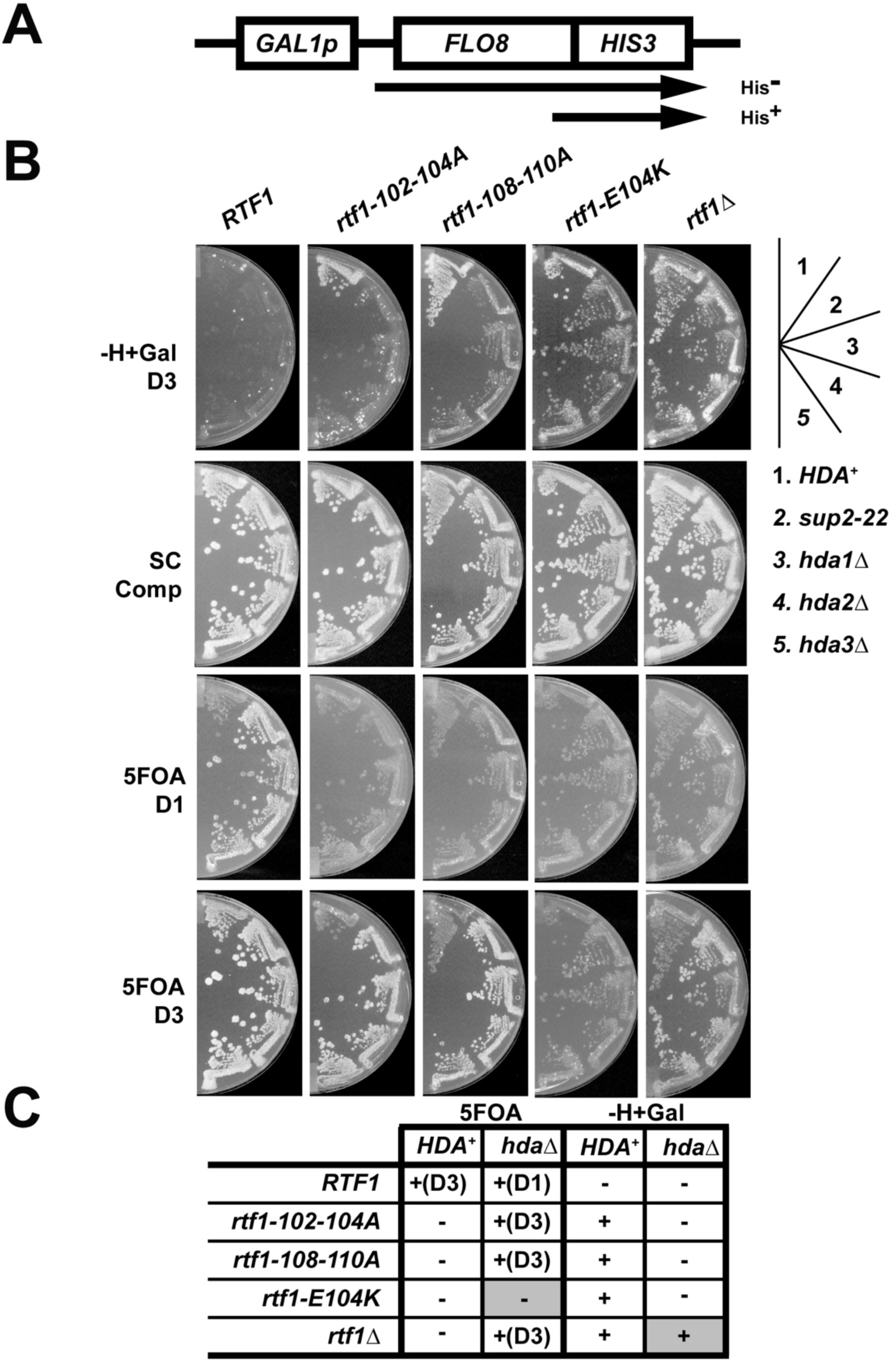
Deletion of any Hda1C subunit suppresses the cryptic initiation phenotype of Rtf1-HMD mutants. Suppression of the cryptic initiation and telomeric silencing defects of *rtf1* point mutations was assessed by replica plating to SC-His+Gal and 5FOA medium, respectively. SC complete (SC Comp) medium was used as a control. Plates were imaged one (D1) or three (D3) days after replica plating and incubating at 30 °C. Growth effects are summarized in the table below the plates. Greyed boxes indicate where the phenotype differed from the other *rtf1* alleles. The following strains were used: KY1523, KY1496, KY2963, KY2934, KY2868, KY2792, KY1519, KY2957, KY2965, KY2898, KY1494, KY1502, KY2973, KY2974, KY2902, KY1526, KY2855, KY2999, KY3001, KY2867, KY1370, KY1520, KY2981, KY2983 and KY2943.

Surprisingly, we identified two strains, with mutations in the same complementation group, in which suppression was not dependent on the presence of a plasmid (see Materials and Methods for details). The chromosomal mutation we chose to analyze, *sup2-22*, suppressed the cryptic initiation phenotype caused by the *rtf1-108-110A* mutation and two other Rtf1 HMD loss-of-function mutations, *rtf1-102-104A* and *rtf1- E104K* (Tomson *et al*. 2011) (Figure 1B, compare sections 1 and 2 in the *rtf1* strains). The *sup2-22* did not suppress the cryptic initiation phenotype of *an rtf1Δ* strain, indicating that suppression is allele-specific.

Bulk segregant analysis followed by whole genome sequencing identified *sup2- 22* as a mutation in the *HDA3* gene, which encodes a non-catalytic subunit of the Hda1C complex (Lee and Kim 2020). This mutation eliminates 283 (out of 655) amino acids from the C-terminus of Hda3. Structural studies indicate that the missing region encompasses a segment necessary for the dimeric interactions between Hda2 and Hda3, which is required for the integrity and function of Hda1C (Wu *et al*. 2001a; Lee *et al*. 2009). We asked whether suppression of the *rtf1* mutations was specific to our original *hda3* mutation or whether loss of any subunit within Hda1C could confer suppression. Complete open reading frame deletions of *HDA1*, *HDA2*, or *HDA3* were created using one-step gene replacement with the *TRP1* gene (Moqtaderi and Geisberg 2013). Removal of any subunit of Hda1C reversed the cryptic initiation phenotype of strains containing *rtf1* point mutations that altered the HMD (Figure 1B, sections 3 to 5); no differences in suppression were detected among the Hda1C mutants. Consistent with results obtained with the *sup2-22* mutation, the cryptic initiation phenotype of strains lacking the entire *RTF1* gene was not suppressed by deletion of individual Hda1C subunits. This result suggests that additional regions of Rtf1 or additional proteins that interact with Rtf1 outside of the HMD are necessary to observe the suppression.

Another phenotype displayed by certain *rtf1* mutants is the inability to silence transcription near telomeres, as assayed using a *TELVR::URA3* reporter (Chu *et al*. 2007; Tomson *et al*. 2011; Van Oss *et al*. 2016). Strains containing this reporter and the wild-type *RTF1* gene grow in the presence of 5FOA, a drug that is toxic to cells expressing *URA3.* Strains, such as *rtf1-108-110A* mutant, that are defective for silencing of the *TELVR::URA3* reporter are 5FOA^S^. Therefore, we used this reporter as a secondary readout for the suppression of the *rtf1* mutations. We originally noted that *RTF1* strains with the *sup2-22* allele grow better on 5FOA-containing media than wild-type strains (Figure 1B, see growth on 5FOA on day one). Consistent with our identification of *sup2-22* as a recessive mutation in *HDA3*, *hda1Δ* mutants were previously shown to have increased resistance to 5FOA, using a telomeric reporter (Rundlett *et al*. 1996; Hang and Smith 2011). The *sup2-22* mutation suppressed the 5FOA sensitivity of two different *rtf1* HMD mutants, *rtf1-102-104A* and *rtf1-108-110A*, as well as the *rtf1Δ* allele. In contrast, *sup2-22* did not suppress the 5FOA sensitivity of a different HMD mutant, *rtf1-E104K*, indicating allele-specific suppression for this phenotype (summarized in Figure 1C).

In addition to effects on telomeric silencing and cryptic initiation from the internal *FLO8* promoter, some mutations that alter the Rtf1 HMD also show a suppressor-of-Ty (Spt^-^) phenotype (Warner *et al*. 2007; Tomson *et al*. 2011; Van Oss *et al*. 2016). At the *his4-912δ* allele, an insertion of a Ty1 δ element in the *HIS4* promoter renders cells unable to grow on media lacking histidine due to an upstream shift in transcription initiation (Winston 1992). However, mutations in many genes regulating transcription and chromatin, including *RTF1*, are able to grow in the absence of histidine because they restore initiation to the native *HIS4* TSS and produce a functional *HIS4* transcript (Stolinski *et al*. 1997). This could be considered another case of cryptic transcription since the native promoter is downstream of the promoter provided by the Ty element (Kaplan *et al*. 2003). Similar to results seen with the *GAL1p-FLO8-HIS3* reporter, the Spt^-^ phenotype of *rtf1-E104K,* but not *rtf1Δ*, was suppressed by deleting *HDA3* (Figure 2).

**Figure 2:**
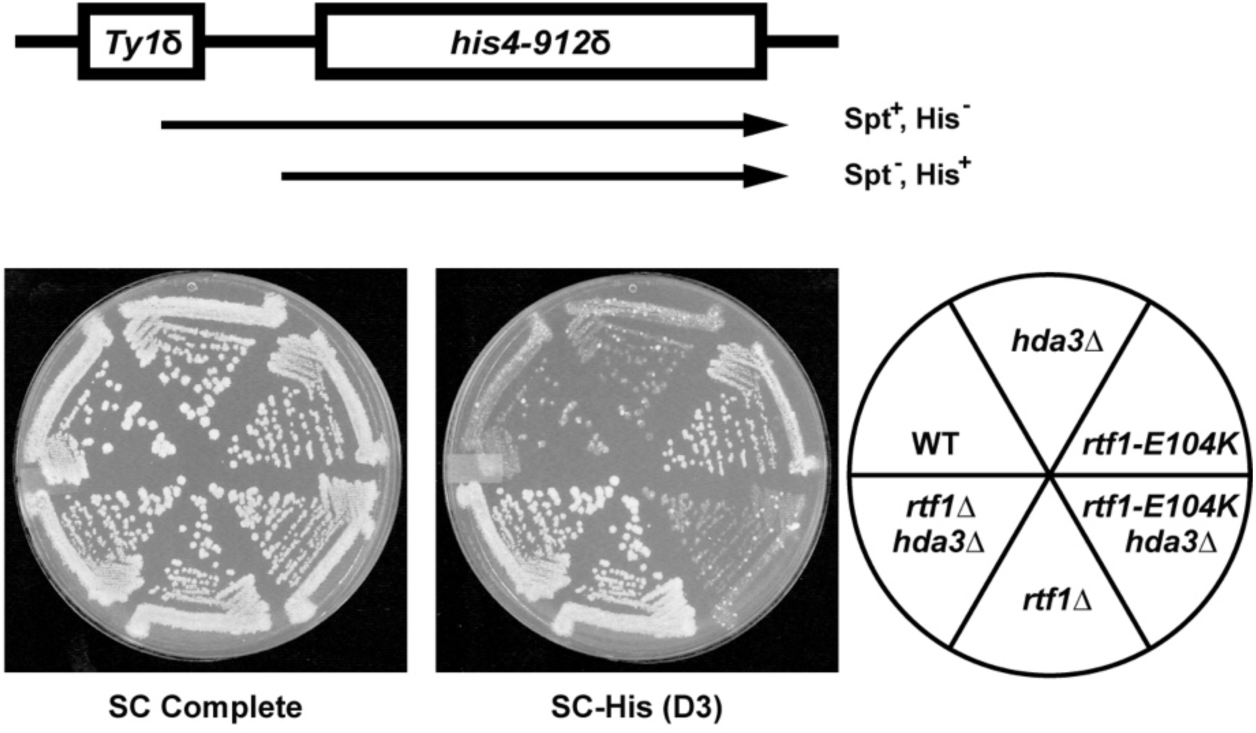
The Spt^-^ phenotype of *rtf1-E104K* is suppressed by *hda3Δ*. The Spt^-^ phenotype of *rtf1-E104K* at the *his4-912δ* locus was detected on medium lacking histidine (SC-His) three days after replica plating; suppression of the phenotype is indicated by reduced growth of the *rtf1-E104K hda3Δ* strain. The following strains were used: KY3191, KY3186, KY1532, KY3178, KY3034 and KY3032.

The identification of loss-of-function mutations in genes encoding an HDAC as suppressors of cryptic initiation was unexpected. Previous work has argued that increased acetylation of histones facilitates cryptic transcription (Rando and Winston 2012; Venkatesh and Workman 2015), likely by relaxing chromatin structure, which could allow spurious pre-initiation complex assembly and the initiation of new transcripts. Thus, mutations that inactivate an HDAC, which would increase histone acetylation, would have been unlikely to reverse the cryptic initiation phenotype of the *rtf1* mutant strains. Due to the unanticipated role of Hda1C identified with the *GAL1p-FLO8-HIS3* reporter, we focused on the coordination between Hda1C and Rtf1 with respect to cryptic initiation.

### Loss of Hda1C does not restore H2Bub in *rtf1* strains

The *rtf1* alleles studied here are unable to stimulate H2Bub *in vivo* (Van Oss *et al*. 2016). One possibility is that suppression of cryptic initiation occurs because loss of Hda1C increases H2Bub levels. However, as shown by western blot analysis, the severe H2Bub defect of the *rtf1* mutants was not rescued by deletion of *HDA1* or *HDA2* (Figure 3A). We also asked if loss of Hda1C recovered H3 methylation marks dependent on a functional Rtf1 HMD and H2Bub (Warner *et al*. 2007; Piro *et al*. 2012; Van Oss *et al*. 2016). While slight changes in global levels of H3K4me2, H3K4me3 and H3K79me2/3 were detected in strains lacking Hda3, we also observed a small increase in total H3 in these strains (Figure S1). When normalized to total H3 levels, the changes in H3K4 or H3K79 methylation marks were less than two-fold, suggesting that suppression of the cryptic initiation phenotypes of the *rtf1* mutants by loss of Hda1C was unlikely due to the recovery of H3K4 or H3K79 methylation.

**Figure 3:**
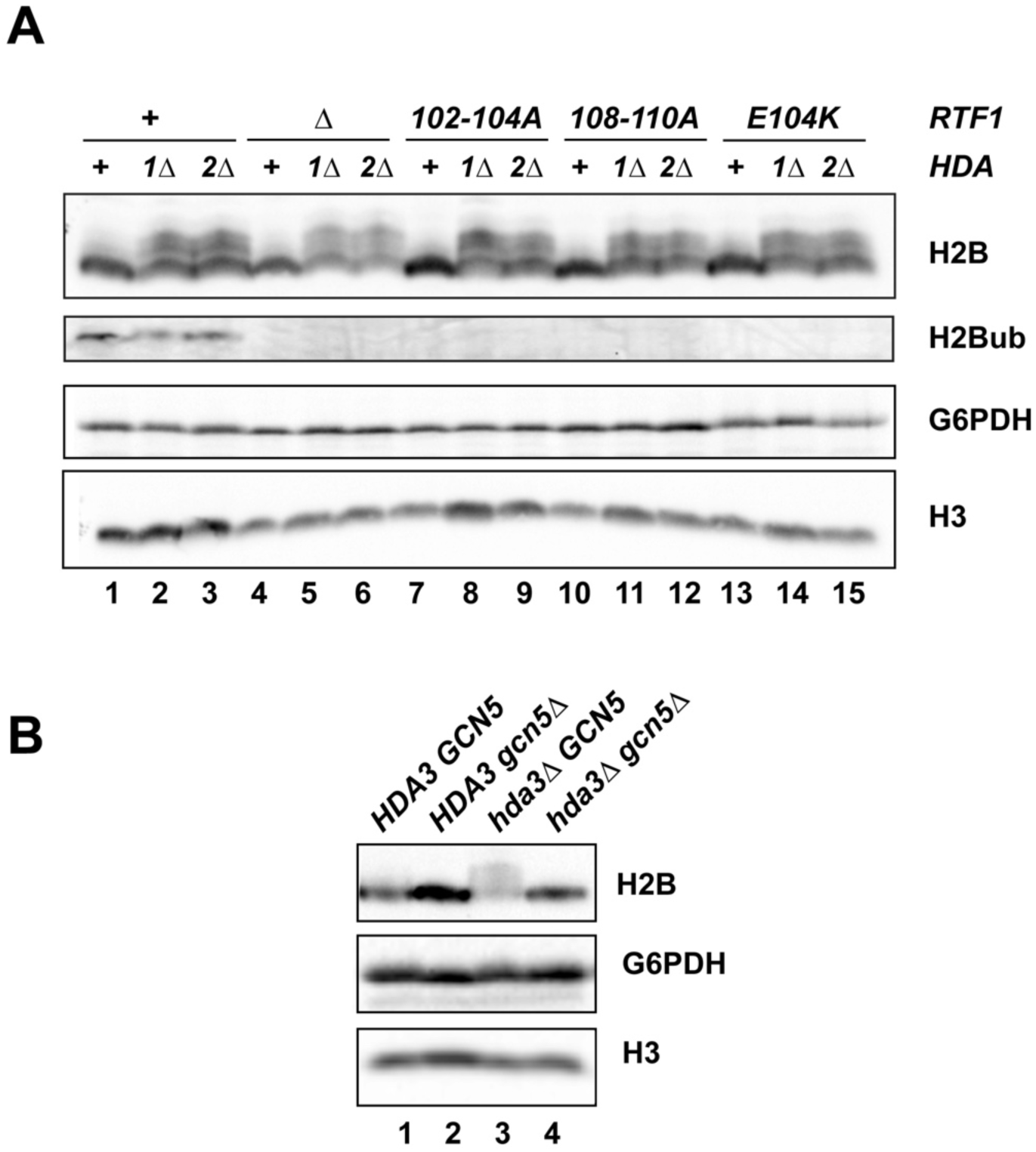
Loss of Hda1C does not restore H2Bub in *rtf1* mutants. (A) Western blot analysis of the indicated *rtf1* mutants in which Hda1C is intact (+) or the *HDA1* (*1Δ*) or *HDA2* (*2Δ*) genes have been deleted. Antibodies against total H2B or H2Bub were used to monitor levels of H2B ubiquitylation. The following strains were used: KY1523, KY2963, KY2934, KY2987, KY2981, KY2983, KY2792, KY2957, KY2965, KY1494, KY2973, KY2974, KY3002, KY2999 and KY3001. (B) Western blot analysis of the indicated strains. The following strains were used: KY433, KY3014, KY2861 and KY3011. In both panels, levels of G6PDH and H3 served as controls.

Strikingly, western blot analysis of total H2B levels showed bands that ran with slower mobility when Hda1C was absent, even in *RTF1* strains (Figure 3A and B). Given the function of Hda1C, we suspected that these bands represented acetylated forms of H2B (Wu *et al*. 2001a, 2001b), stabilized in the absence of the deacetylase complex. As the Gcn5 HAT was previously shown to contribute to H2B acetylation (Grant *et al*. 1997; Suka *et al*. 2001), we asked if the slower migrating forms of H2B were dependent on Gcn5. Indeed, in *gcn5Δ hda3Δ* strains, the slower migrating anti- H2B-reactive bands were no longer observed (Figure 3B, lane 4), suggesting that these bands represent acetylated forms of H2B which are elevated in the absence of Hda1C.

### Loss of H2Bub leads to Hda1C-dependent cryptic initiation

Since the *HDA* mutations suppress the cryptic initiation phenotype of *rtf1* mutants that are severely defective in H2Bub, we assessed the cryptic initiation phenotype of other strains lacking H2Bub and the dependence on Hda1C for this phenotype. We analyzed strains expressing an H2B mutant protein that cannot be ubiquitylated, H2B-K123R, as the only form of H2B and a strain lacking Rad6, the ubiquitin conjugase that targets H2B K123. In both cases, when *HDA3* is wild type, we observed growth on SC-His+Gal medium, indicating cryptic initiation from the *GAL1p-FLO8-HIS3* reporter (Figure 4A and B), identifying a role for H2B K123 in controlling cryptic initiation using this reporter. Consistent with the suppression of *rtf1* mutations that are strongly defective in promoting H2Bub, *hda3Δ* also suppresses the cryptic initiation phenotype of *htb1- K123R* and *rad6Δ* strains (Figure 4A and B). Together these results suggest that the role of Hda1C in supporting cryptic initiation in the *rtf1* mutants occurs downstream of the effects of losing H2Bub.

**Figure 4:**
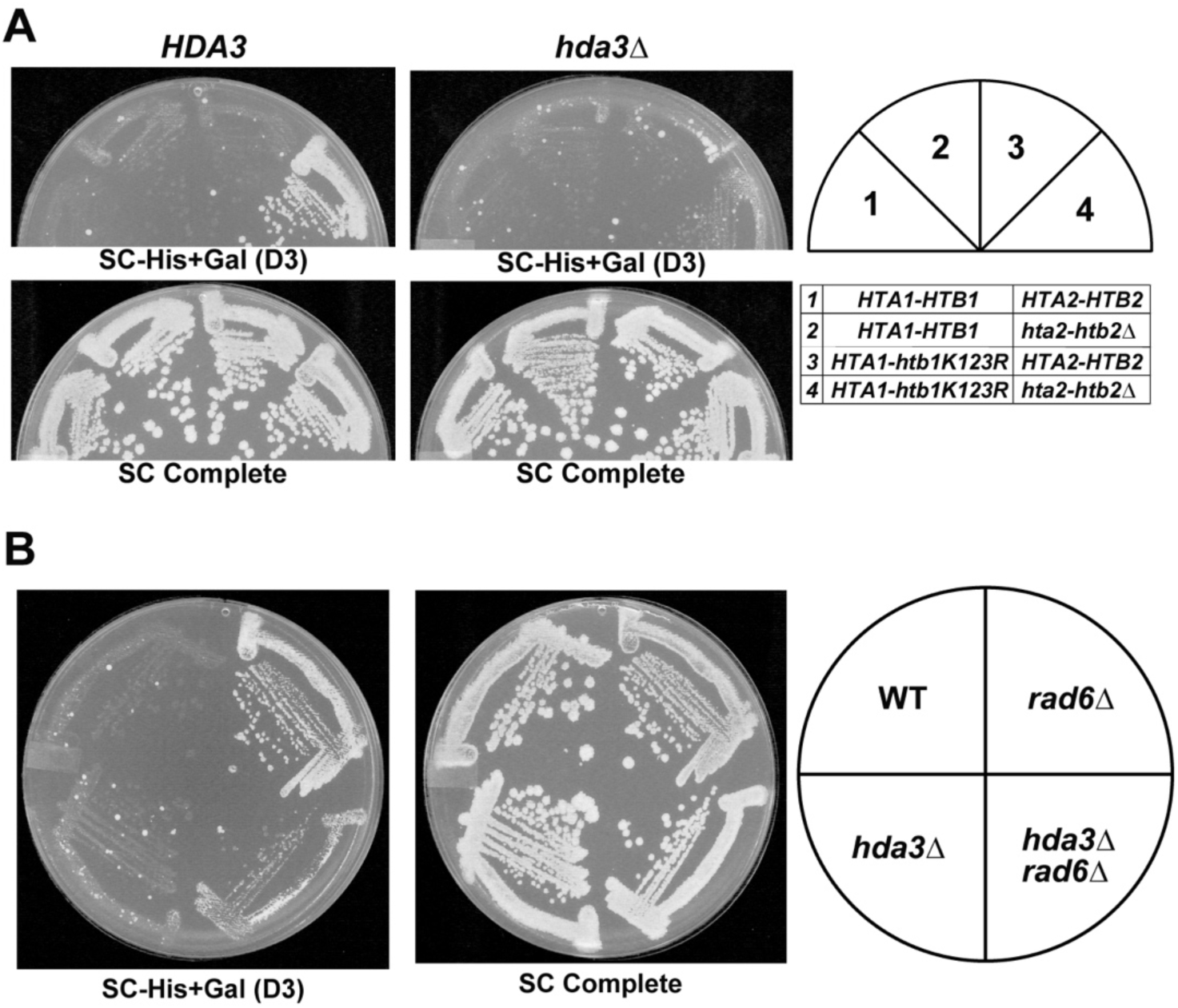
Deletion of *HDA3* suppresses defects in the H2Bub pathway. (A) A strain carrying the H2B-K123R substitution as the only source of H2B (section 4) exhibits cryptic initiation when Hda3 is present (left), but not when *HDA3* has been deleted (right). The following strains were used: KY1523, KY2878, KY2929, KY2876, KY2868, KY2921, KY2924 and KY2910. (B) A strain in which *RAD6* has been deleted exhibits cryptic initiation when Hda3 is present (top right quadrant), but not when *HDA3* has been deleted (bottom right quadrant). The following strains were used: KY1523, KY2832, KY2918 and KY2868. For both panels, plates were imaged three days after replica plating and incubation at 30 °C.

### The genetic interactions between Rtf1 and Hda1C are distinct from those involving other HDACs and chromatin regulators

Contrary to our results with Hda1C, previous studies showed that other HDACs suppress cryptic initiation; that is, in their absence, increased levels of cryptic transcription are detected (Cheung *et al*. 2008; Silva *et al*. 2012). Given our unexpected result, we asked whether mutations in genes encoding other HDACs, in particular, Rpd3(S) (*rco1Δ*) (Carrozza *et al*. 2005; Keogh *et al*. 2005) or Set3C (*set3Δ*) (Pijnappel *et al*. 2001; Kim *et al*. 2012), would also suppress the transcription defect in *rtf1-102-104A* strains. By using a deletion of *RCO1,* we specifically targeted the Rpd3(S) complex, which has been previously connected to regulation of intragenic cryptic transcription. As expected (Carrozza *et al*. 2005; Cheung *et al*. 2008; Silva *et al*. 2012; McDaniel *et al*. 2016), a strain lacking the Rpd3S subunit Rco1 and containing the *GAL1p-FLO8-HIS3* reporter grew on Sc-His+Gal media (Figure 5, Sc-His+Gal (D3)). In fact, the double mutant, *rco1Δ rtf1-102-104A*, grew better than either single mutant (Figure 5, Sc-His+Gal (D2)), suggesting the mutations act in different pathways to affect cryptic initiation. In addition, *rpd3Δ* strains, lacking both the Rpd3(S) and Rpd3(L) complexes, showed similar results (Figure S2), though strains lacking Rpd3 grow more slowly. Deletion of *SET3*, which eliminates activity from both the Hos2 and Hst2 histone deacetylases in Set3C, did not lead to a cryptic initiation phenotype as measured by the *GAL1p-FLO8-HIS3* reporter, in agreement with previous reports that *set3Δ* did not elicit transcription from the cryptic promoters in the native *FLO8* or *STE11* genes (Kim and Buratowski 2009). Furthermore, unlike *hda3Δ*, *set3Δ* did not suppress the cryptic initiation phenotype of *rtf1-102-104A*. Therefore, suppression of cryptic transcription in this *rtf1* mutant by *hda3Δ* is not a common feature of HDAC mutants.

**Figure 5:**
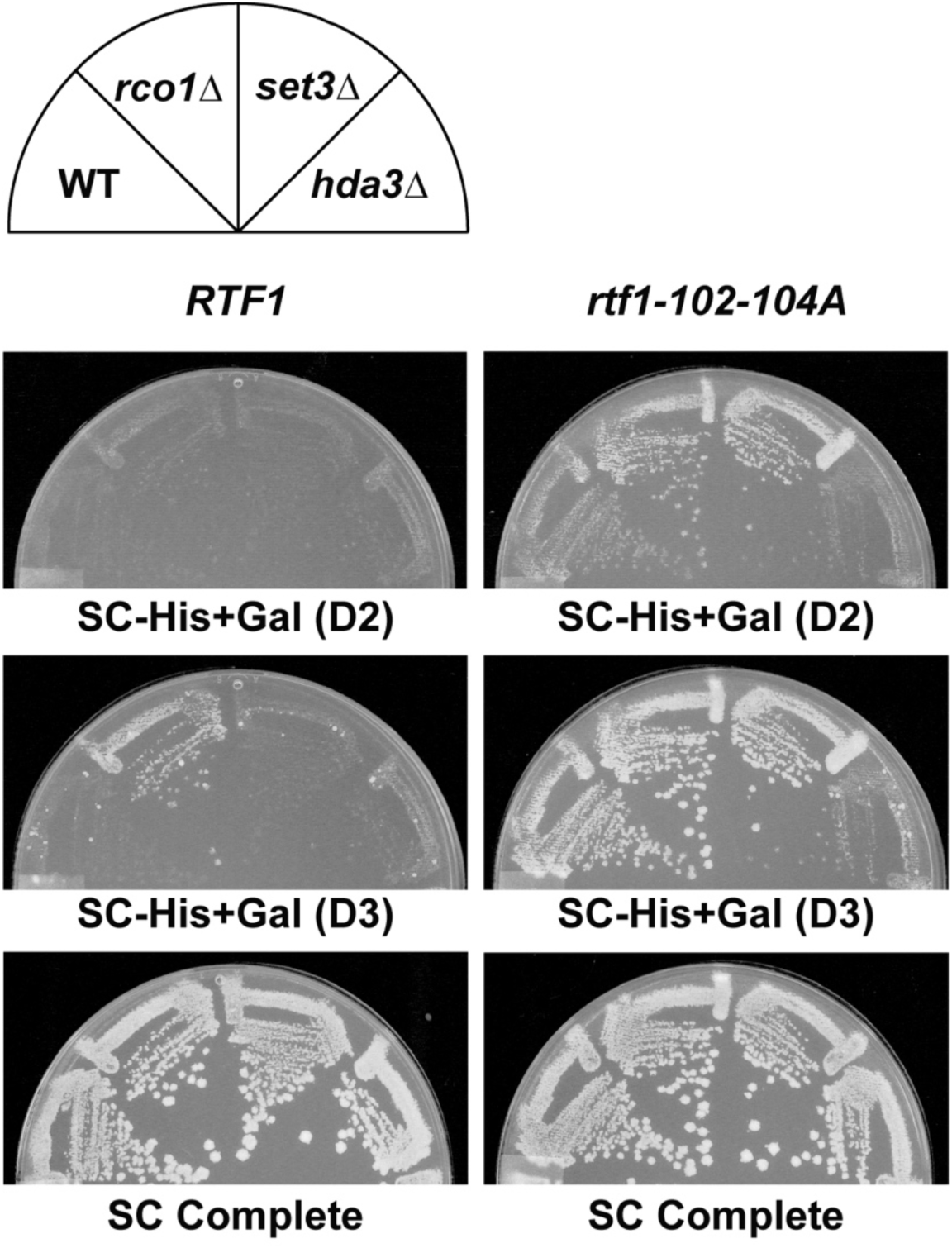
Mutations that disrupt other HDACs do not suppress the cryptic initiation phenotype of an *rtf1* HMD mutant. Strains mutated for the Rpd3(S) (*rco1Δ*) or Set3C (*set3Δ*) HDACs and containing either *RTF1* or *rtf1-102-104A* were replica plated to SC-His+Gal or SC Complete media and scanned after two (D2) or three (D3) days of incubation at 30 °C. The following strains were used: KY1523, KY3057, KY3042, KY2868, KY2792, KY3053, KY3038 and KY2898.

Next, we investigated if the effect of Hda1C was specific to strains defective for H2Bub. Many strains with mutations in genes encoding chromatin and transcription factors show cryptic initiation on the *FLO8-HIS3* reporter (Cheung *et al*. 2008; Quan and Hartzog 2010; Silva *et al*. 2012). These include strains lacking or mutated in the H3K36 methyltransferase Set2, the chromatin remodeling enzyme Chd1, and the histone chaperones Spt6 and Spt16. In contrast to *rtf1* mutants, *hda3Δ* did not suppress the cryptic initiation phenotypes of strains lacking Set2 or Chd1 or containing mutations in *SPT6* or *SPT16* (Figure 6A and B). In fact, *set2Δ hda3Δ* and *spt16-197 hda3Δ* double mutants grew better on the Sc-His+Gal medium than *set2Δ* or *spt16-197* single mutant strains. These results indicate that loss of Hda1C is not simply suppressing cryptic initiation by reducing transcription from the *GAL1* promoter and that the genetic interaction we identified between genes affecting H2Bub and Hda1C is distinct among genes that affect cryptic initiation.

**Figure 6:**
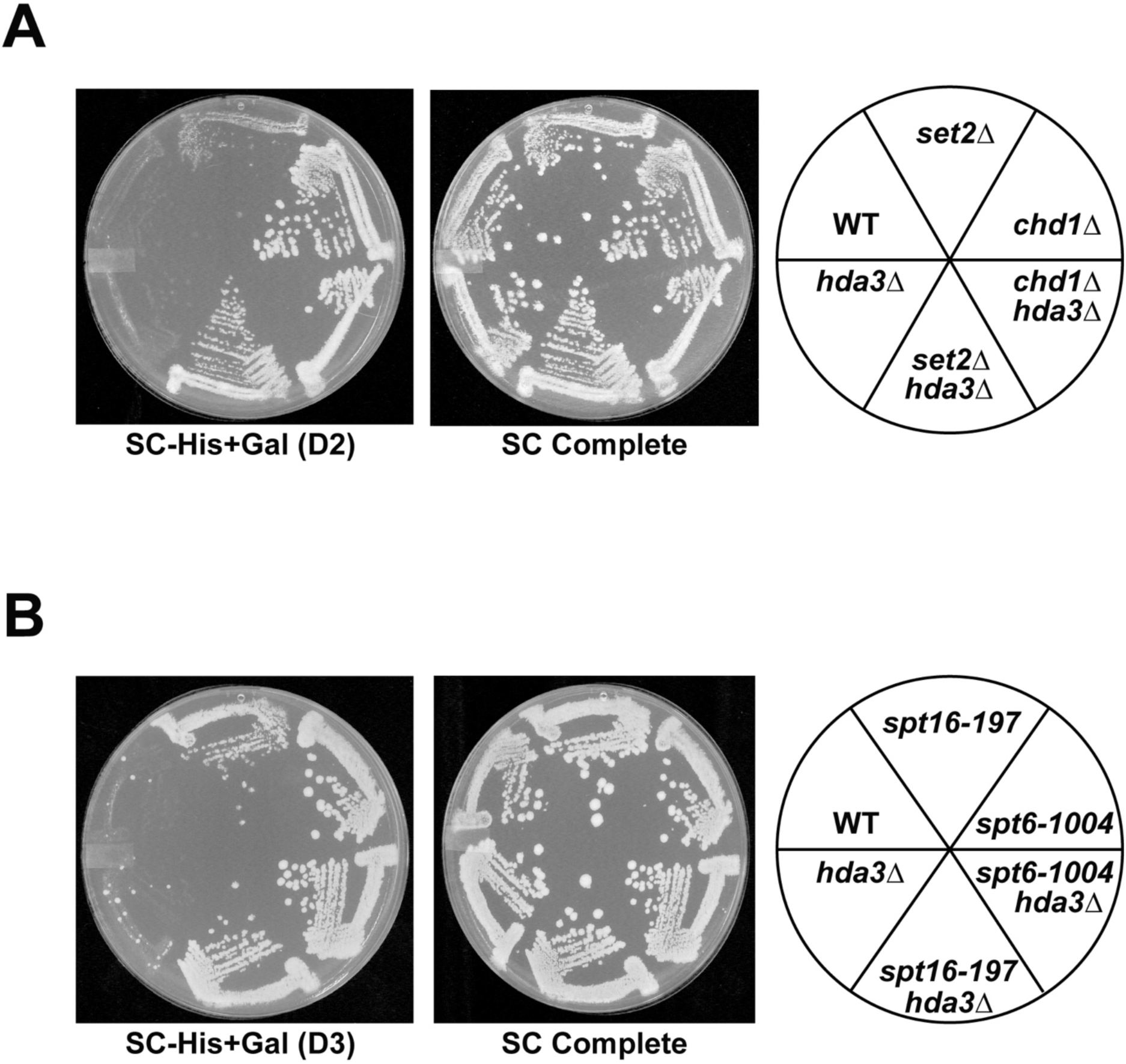
Deletion of *HDA3* does not suppress the cryptic initiation phenotypes of strains defective in other chromatin regulators. Strains containing the indicated genotypes were replica plated to SC-His+Gal or SC Complete media and scanned after two (D2) or three (D3) days of incubation at 30 °C. The following strains were used: (A) KY1523, KY3200, KY3198, KY3199, KY3201 and KY2868 and (B) KY1523, KY3073, KY3084, KY3082, KY3093 and KY2868.

### Sas3 and H3K14 suppress cryptic initiation in an *rtf1* mutant

Since loss of Hda1C suppresses the cryptic initiation phenotype of *rtf1* mutants, we hypothesized that deletion of a HAT that works in opposition to Hda1C might exacerbate the cryptic initiation phenotype of the *rtf1* mutants and reverse the effect of inactivating Hda1C. Based on our western blot results showing that deletion of *GCN5* reversed the H2B mobility shift caused by *hda3Δ*, we investigated if Gcn5 was the relevant HAT. Gcn5 is the catalytic subunit of several HAT complexes (Lee and Workman 2007), any of which might be involved in regulating cryptic initiation. If Gcn5 antagonized Hda1C, then the *rtf1-102-104A hda3Δ gcn5Δ* triple mutant would be expected to phenocopy the *rtf1-102-104A* single mutant and grow on Sc-His+Gal medium. Instead, the triple mutant was strongly His^-^ (Figure 7, right panel, section 4). In addition, on its own, the *gcn5Δ* mutation suppressed the cryptic initiation phenotype of the *rtf1-102-104A* mutation (Figure 7, right panel, section 3). These results suggest that suppression of *rtf1-102-104A* by *hda3Δ* is not easily explained by increased acetylation of a Gcn5-dependent target protein.

**Figure 7:**
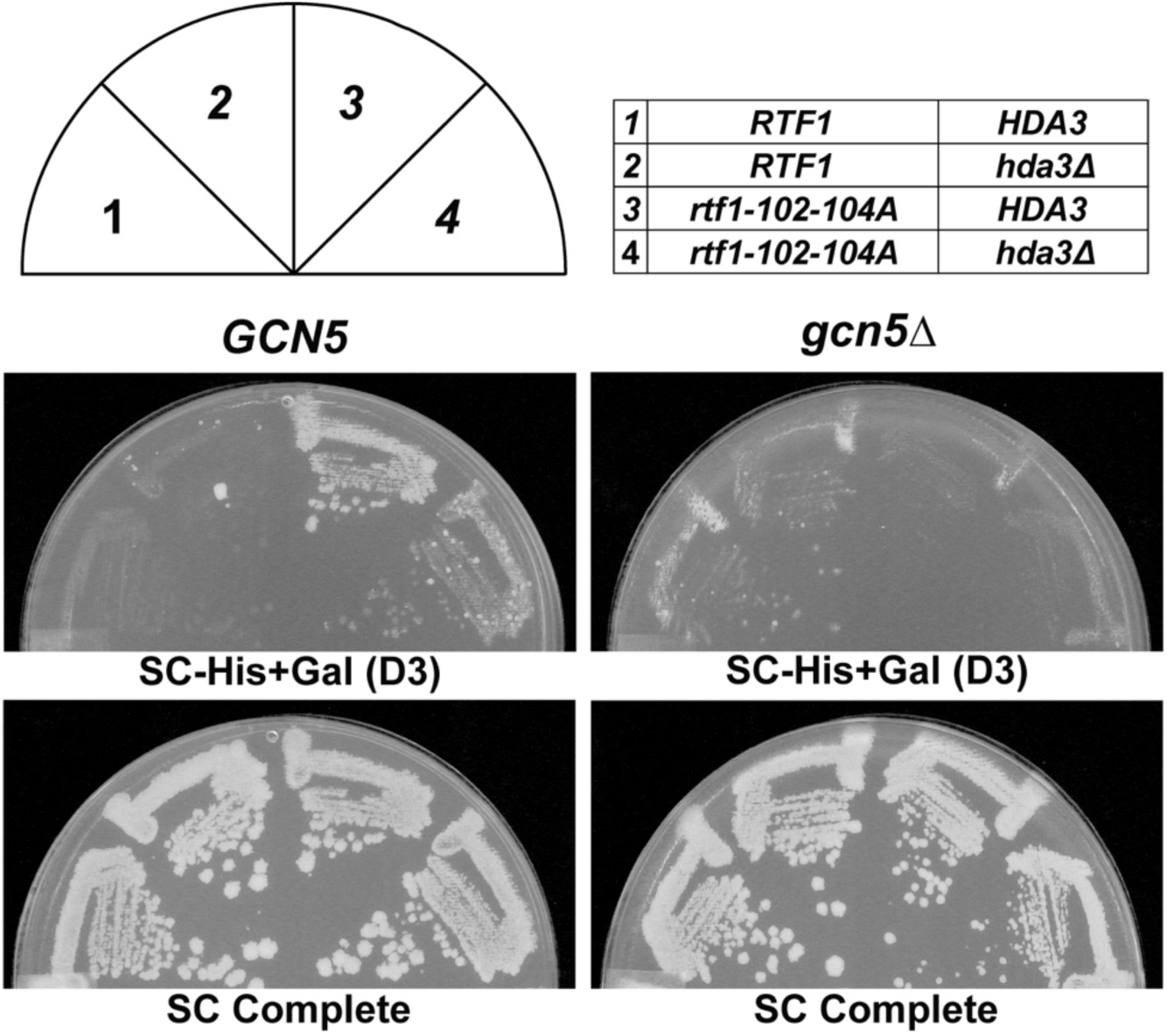
Deletion of *GCN5* suppresses the cryptic initiation phenotype of an *rtf1* HMD mutant. Left panels: *GCN5* strains containing *hda3* and *rtf1* mutations as indicated in the table (top right). Right panels: *gcn5Δ* strains containing *hda3* and *rtf1* mutations as indicated in the table (top right). Plates were scanned three days after replica plating to the indicated media. The following strains were used: KY1523, KY2868, KY2792, KY2898,.KY3117, KY3115, KY3114 and KY3106.

Because the *rtf1* HMD mutants exhibit telomeric silencing defects and Hda1C also plays a role in silencing, we asked whether deletion of two HATs previously implicated in transcriptional silencing would affect the cryptic initiation phenotype of the *rtf1* mutants. *SAS2* was first discovered in a screen for genes affecting transcriptional silencing at the mating-type locus and *SAS3* was subsequently uncovered by its similarity to *SAS2* (Reifsnyder *et al*. 1996). Sas2 and Sas3 are the catalytic subunits of the HATs SAS and NuA3, respectively (John *et al*. 2000; Shia *et al*. 2005). While deletion of *SAS2* had little effect, deletion of *SAS3* exacerbated the cryptic initiation phenotype of the *rtf1* mutants, as indicated by enhanced growth on Sc-His+Gal medium. Interestingly, *sas3Δ* on its own did not confer a cryptic initiation phenotype (Figure 8A). Notably, Sas3 and Hda3 appear to oppose each other because deleting both proteins in the context of *rtf1-108-110A* gave a cryptic initiation phenotype nearly identical to the *rtf1* allele alone (Figure 8B). These results suggest that the acetylation state of one or more shared targets of Hda1C and Sas3 modulates the cryptic initiation phenotype of the *rtf1* mutants.

**Figure 8:**
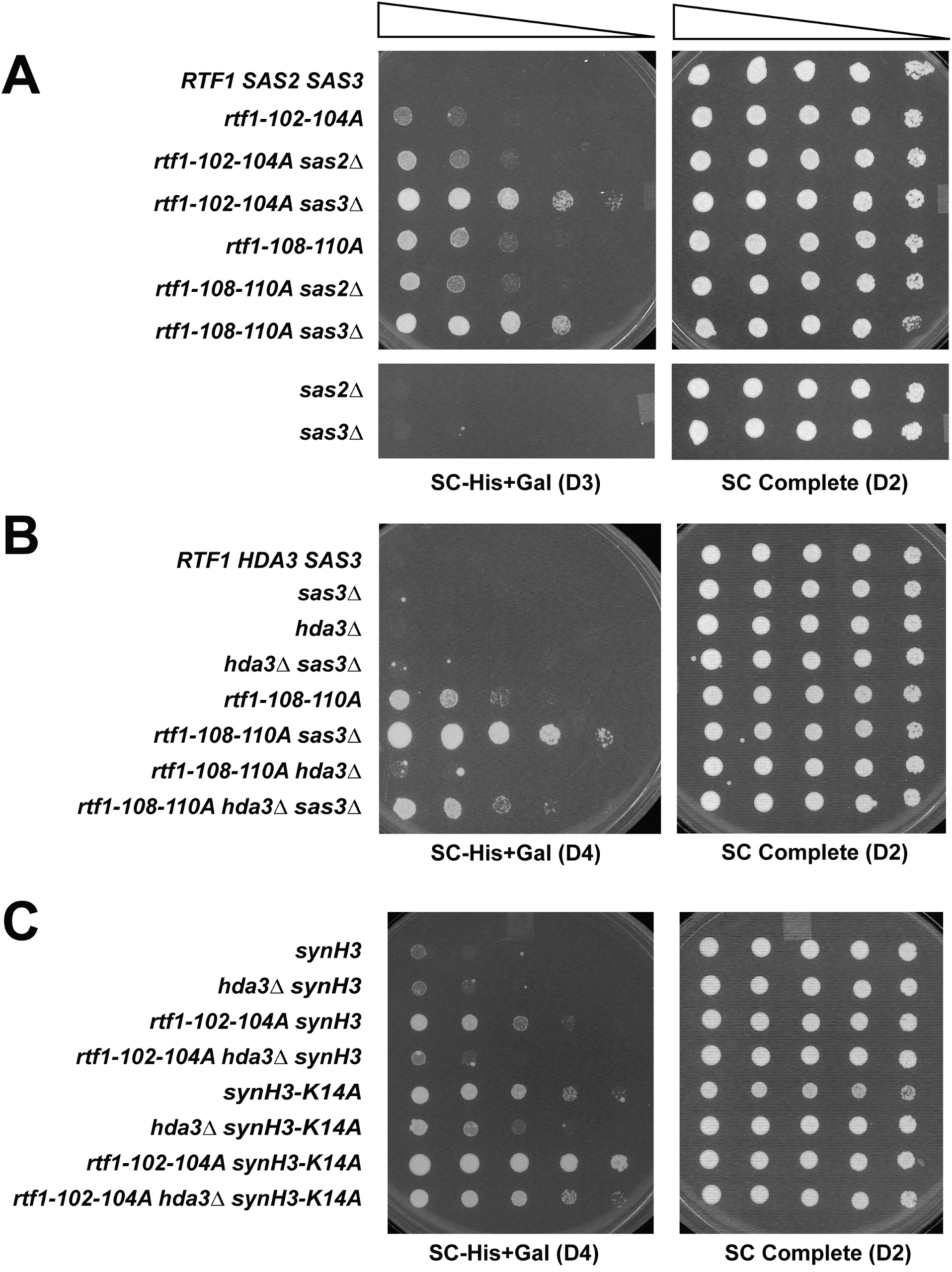
Mutation of *SAS3* or H3K14 enhances the cryptic initiation phenotype of *rtf1* HMD mutants. (A,B) Strains with the indicated genotypes were grown as described in the Materials and Methods and serial dilutions were spotted onto SC-His+Gal and SC Complete media and allowed to grow for two (D2), three (D3) or four (D4) days at 30 °C before imaging. (C) Strains deleted for *HHT1-HHF1* and carrying a synthetic copy of *HHT2-HHF2* (synH3), either wild-type or H3K14A, were used in genetic crosses to create double and triple mutants with *rtf1-102-104K* and *hda3Δ*. Strains were grown on the indicated media for two to four days at 30 °C. The following strains were used: (A) KY1491, KY2792, KY3366, KY3371, KY1490, KY3369, KY3372, KY3362 and KY3364; (B) KY3423, KY3431, KY3427, KY3438, KY3425, KY3434, KY3429 and KY3441; (C) KA255, KA256, KA257, KA258, KA295, KA296, KA297 and KA298

One target of Sas3 is H3K14 (Taverna *et al*. 2006). We took advantage of an integrated synthetic histone mutant library (Dai *et al*. 2008) to create strains in which the only copy of H3 had a lysine-to-alanine substitution at position 14 (H3K14A). Unlike *sas3Δ* strains, the H3K14A mutant showed cryptic initiation using the *FLO8-HIS3* reporter (Figure 8C). The distinct phenotypes of the *sas3Δ* and H3K14A mutants may be due to residual acetylation of H3K14 in the *sas3Δ* strain, as previous studies showed that H3K14 is also acetylated by Gcn5 (Church *et al*. 2017). However, like the *sas3Δ* mutation, the H3K14A substitution strongly enhanced the cryptic initiation phenotype of the *rtf1-102-104A* mutant. Deleting *HDA3* only partially reversed the *rtf1-102-104A H3K14A* cryptic initiation phenotype. Unlike the *rtf1-102-104A hda3Δ synH3* control strain, which carries a synthetic version of wild-type H3, the *rtf1-102-104A hda3Δ synH3-K14A* remained His^+^. Together, these results suggest that Sas3 represses the cryptic initiation phenotype of *rtf1* mutants and Hda1C opposes Sas3 in this process. One possible target for their action is H3K14. However, because *hda3Δ* partially suppresses the cryptic initiation phenotype of the synH3-K14A mutant, other Hda1C targets that control chromatin accessibility must exist.

## DISCUSSION

In this study we discovered an unexpected connection between Rtf1, a component of the Paf1 transcription elongation complex necessary for H2Bub, and Hda1C, a histone deacetylase complex, in controlling cryptic transcription initiation within a gene body. Previous studies have shown that deletion of certain HDACs, including Rpd3(S) or Set3C, causes cryptic initiation (Carrozza *et al*. 2005; Keogh *et al*. 2005; Li *et al*. 2007a, 2007b; Kim *et al*. 2012). This is consistent with the expectation that increased histone acetylation promotes a permissive chromatin environment. In contrast, our study shows that loss of Hda1C suppresses the cryptic initiation phenotype of strains deficient in or completely lacking in H2Bub. The interplay between H2Bub and Hda1C in controlling cryptic transcription initiation is specific, because deletion of *HDA3* did not reduce cryptic initiation in strains lacking several other chromatin-related proteins and mutations in genes encoding other HDACs did not suppress the cryptic initiation phenotype of an *rtf1* mutant defective in H2Bub.

We and others have shown that mutations in *RTF1* and genes encoding other proteins required for H2Bub cause cryptic initiation within the *FLO8* gene (Cheung *et al*. 2008; Fleming *et al*. 2008; Tomson *et al*. 2011; Silva *et al*. 2012). While H2Bub is required for H3K4me and H3K79me *in vivo*, previous studies found that strains lacking Set1 or Dot1, the methyltransferases responsible for H3K4me and H3K79me, respectively, did not exhibit cryptic initiation at the *FLO8-HIS3* reporter (Quan and Hartzog 2010). This suggests that H2Bub functions independently of these downstream H3K4 and H3K79 modifications in preventing cryptic initiation. H2Bub has been shown to stabilize nucleosomes, particularly in the middle and ends of genes (Chandrasekharan *et al*. 2009; Batta *et al*. 2011). Recent work using optical tweezers has confirmed the increase in stability of nucleosomes containing H2Bub, and this increase in stability is an energetic barrier to passage by Pol II (Chen *et al*. 2019). One possibility based on our findings is that Hda1C may contribute to the instability of nucleosomes lacking H2Bub, most likely by deacetylating a histone or other target.

Interestingly, deletion of Hda1C did not suppress the cryptic initiation phenotype of strains completely lacking Rtf1. This suggests that the absence of other functional domains within Rtf1, outside of the HMD, contributes to cryptic initiation in a way that is not suppressed by mutations in genes encoding Hda1C. In addition to the HMD, Rtf1 has separable domains important for its interactions with the Pol II elongation complex, Chd1, and other Paf1C subunits (Simic *et al*. 2003; Warner *et al*. 2007; Mayekar *et al*. 2013; Wier *et al*. 2013; Vos *et al*. 2020). The N-terminus of Rtf1 is important for the recruitment of Chd1 to open reading frames (Warner *et al*. 2007), and we found that deletion of *CHD1* causes a cryptic initiation phenotype that cannot be suppressed by *hda3Δ*. This observation suggests that Chd1 and Hda1C affect cryptic initiation through different pathways or that the effect of Chd1 is upstream of the effect of Hda1C. Therefore, while a complete deletion of Rtf1 can affect transcription and chromatin through several mechanisms, our data suggest that only the effect on H2Bub is suppressed by loss of Hda1C.

What is the target of Hda1C that creates a permissive condition for initiation of transcription within the coding sequence? Cells lacking Hda1C have increased acetylation at several positions on H2B, H3 and H4, including H3K14 (Carmen *et al*. 1996; Rundlett *et al*. 1996; Wu *et al*. 2001a, 2001b; Yang and Seto 2008; Islam *et al*. 2011; Ha *et al*. 2019; Lee and Kim 2020). H3K14Ac has been previously shown to help maintain proper levels of H3K4me3 by inhibiting the Jhd2 histone demethylase (Maltby *et al*. 2012). Furthermore, in a positive feedback loop, methylation of H3K4, a modification dependent on H2Bub, promotes H3K14 acetylation by Sas3 as part of the NuA3 HAT complex, through the binding of the Yng1 subunit of NuA3 (Taverna *et al*. 2006; Martin *et al*. 2017). Our results demonstrate that, in the context of an *rtf1* mutation, the absence of Sas3 or H3K14 enhances cryptic initiation, supporting the idea that an appropriate level or pattern of histone acetylation marks is needed to prevent cryptic initiation. While Hda1C and Sas3 appear to work in opposition in controlling cryptic initiation, the role of Hda3 in promoting cryptic initiation in *rtf1* mutants cannot be solely through deacetylation of H3K14, as we observe partial suppression of the cryptic initiation of H3K14A by *hda3Δ*.

The enrichment of Hda1C on the bodies of active genes and its interactions with RNA and Pol II argue for a role in controlling histone acetylation levels during transcription elongation (Govind *et al*. 2010; Ha *et al*. 2019; Lee and Kim 2020). Moreover, a recent study showed that Hda1C reduces H4 acetylation within the coding regions of genes undergoing high levels of transcription (Ha *et al*. 2019; Lee and Kim 2020). With the *FLO8-HIS3* reporter used here, high levels of transcription driven by the inducible *GAL1* promoter are required to detect cryptic initiation in the *rtf1* HMD mutants (data not shown; see also (Silva *et al*. 2012)). Surprisingly, by a mechanism not understood, *hda1Δ* cells, with higher levels of H4 acetylation, exhibit higher histone occupancy levels, especially on longer genes where the effect of *hda1Δ* on H4 acetylation is more evident (Ha *et al*. 2019). Increasing histone occupancy within gene bodies could be one mechanism by which loss of Hda1C counteracts the disruption in chromatin structure caused by loss of H2Bub.

In summary, our study has uncovered an interplay between the H2Bub pathway and Hda1C in controlling chromatin accessibility during transcription elongation. Unexpectedly, a functional Hda1C permits cryptic initiation in cells lacking H2Bub. While cryptic initiation is most frequently observed under mutant conditions, it is possible that the dynamic removal of H2Bub during transcription elongation could give rise to cryptic transcripts in wild-type cells, albeit at low levels. In this scenario, the activity of Hda1C could regulate the output of these cryptic transcripts. Whether these transcripts would have biological consequences, as has been reported for some internally initiated transcripts (McKnight *et al*. 2014; Tamarkin-Ben-Harush *et al*. 2017), is unclear. Future work will be needed to elucidate the target(s) through which Hda1C impacts cryptic initiation in opposition to H2Bub. Recent technical and computational advances in detecting transcript initiation events within genes bodies (Pelechano *et al*. 2013; Wei *et al*. 2019) offer the opportunity to study this and additional examples of crosstalk between epigenetic regulators in maintaining chromatin structure during transcription elongation.

## ACKNOWLEDGMENTS

We are grateful to Kristin Klucevsek for creating the *sas2Δ* and *sas3Δ* strains, Daniel Sheidy for technical assistance in performing the genetic screen for *rtf1* suppressors, Graham Hatfull and members of his research group for assistance with genomic sequencing, and members of the Arndt, Hainer, Kaplan and Lee laboratories for their insightful comments. This research was supported in part by the University of Pittsburgh Center for Research Computing through the resources provided and by the following funding sources: National Institutes of Health (NIH) grant R01GM052593 to K.M.A., NIH predoctoral fellowship (F31GM129917) to M.A.E., and a Colella Research Fellowship to R.A.K.

---

**Figure S1.**
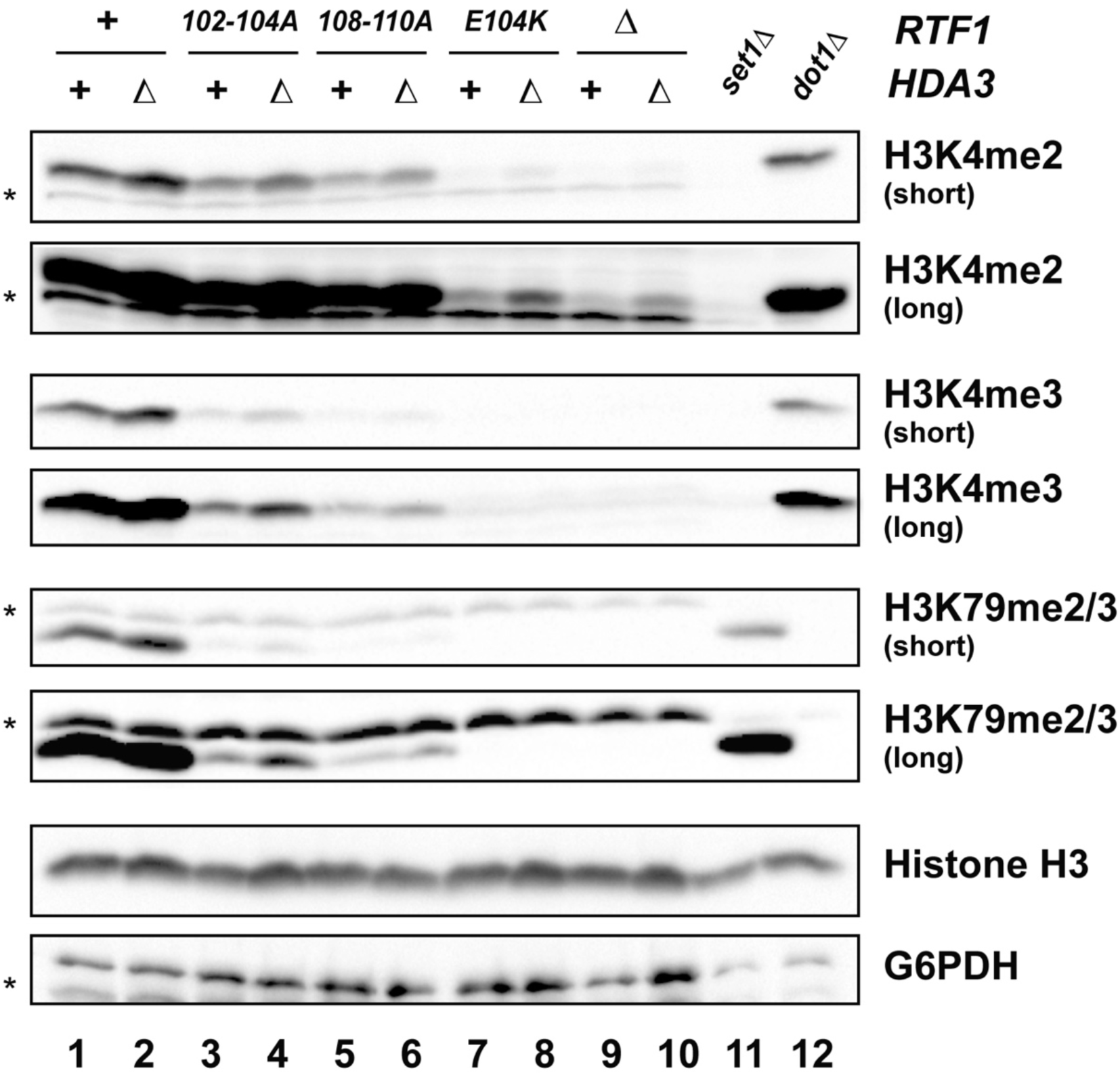
Strains lacking Hda3 show a slight increase in H3K4 and H3K79 methylation. Western blot analysis of the indicated *rtf1* mutants containing or lacking *HDA3*. For blots using antibodies against histone methylation marks, two exposures are shown to visualize the effects in the *rtf1* mutants; in the shorter exposures, no bands are overexposed (saturated). Lanes 11 and 12 show extracts from control strains that lack the histone methyltransferases for H3K4 (*set1Δ*) and H3K79 (*dot1Δ*). Asterisks (*) indicate background bands associated with the non-specific hybridization of the antibodies. The following strains were used: KY1523, KY2868, KY2792, KY2898, KY1494, KY2902, KY3002, KY2867, KY2987, KY2943, KY1715 and KY1717.

**Figure S2:**
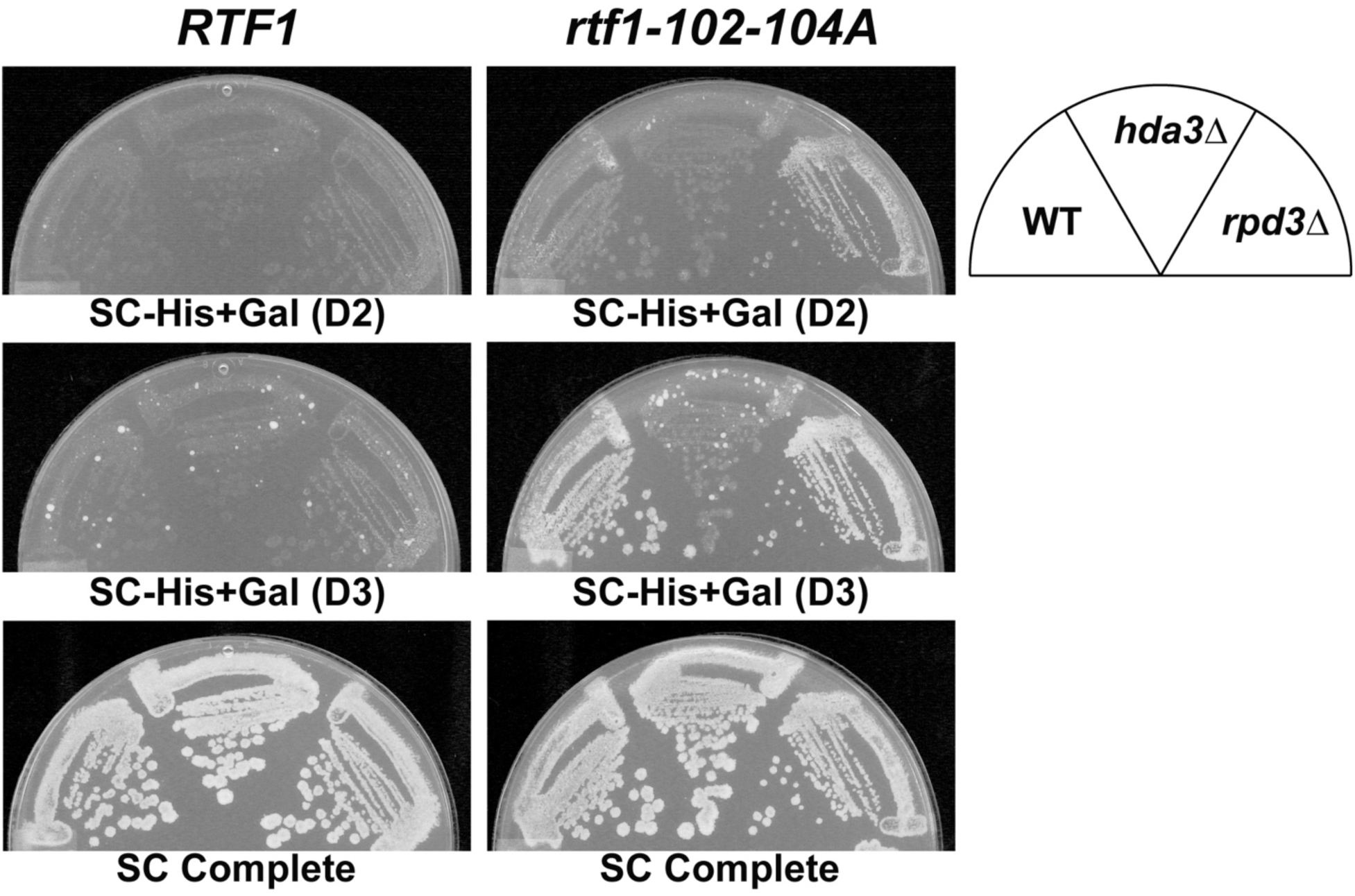
Deletion of *RPD3* does not suppress the cryptic initiation phenotype of an Rtf1 HMD mutant. Strains with the indicated genotypes were replica plated to SC-His+Gal or SC Complete media and scanned after two (D2) or three (D3) days of incubation at 30 °C. The following strains were used: KY1523, KY2868, KY3070, KY2792, KY2898 and KY3063.

**Table S1.**
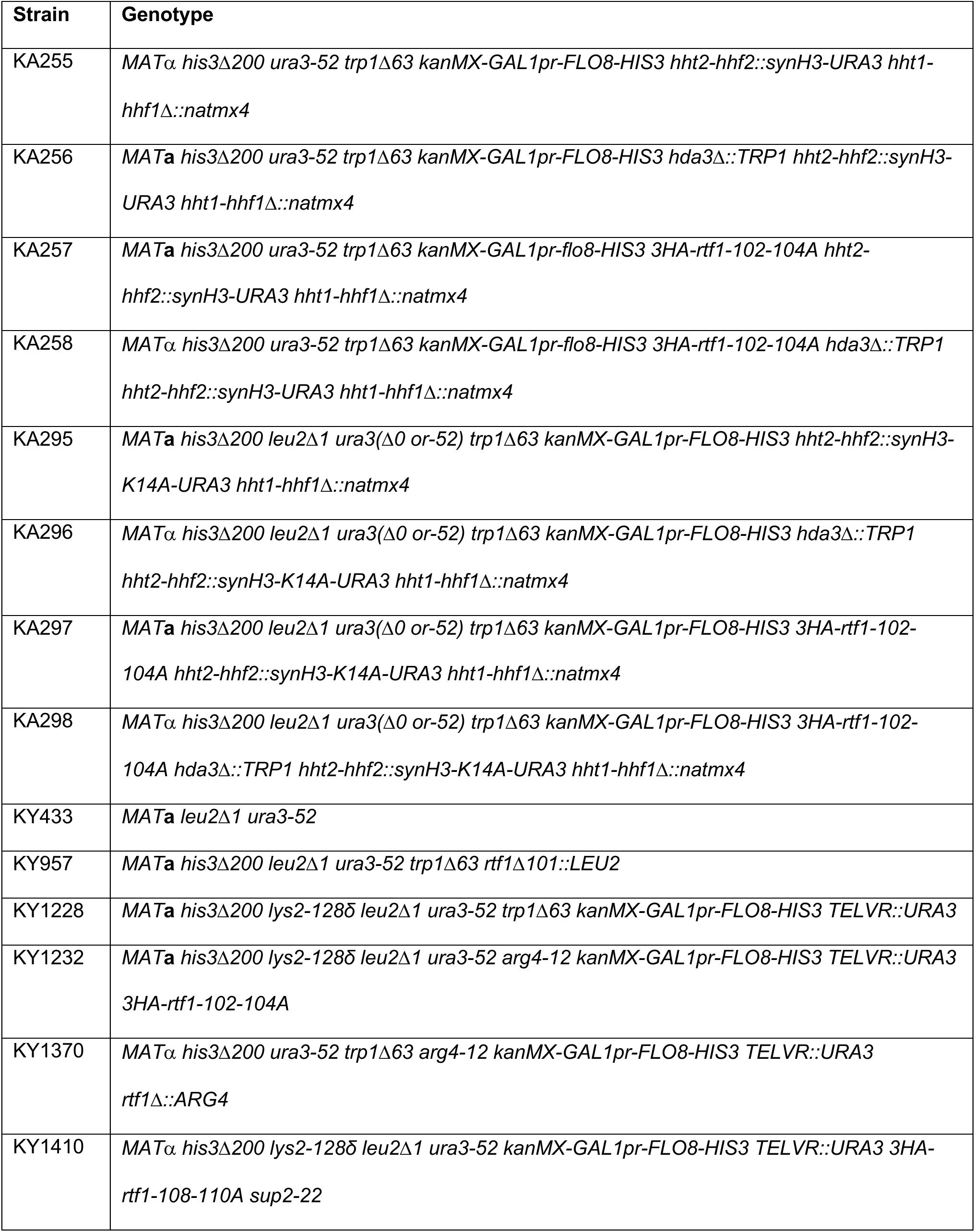
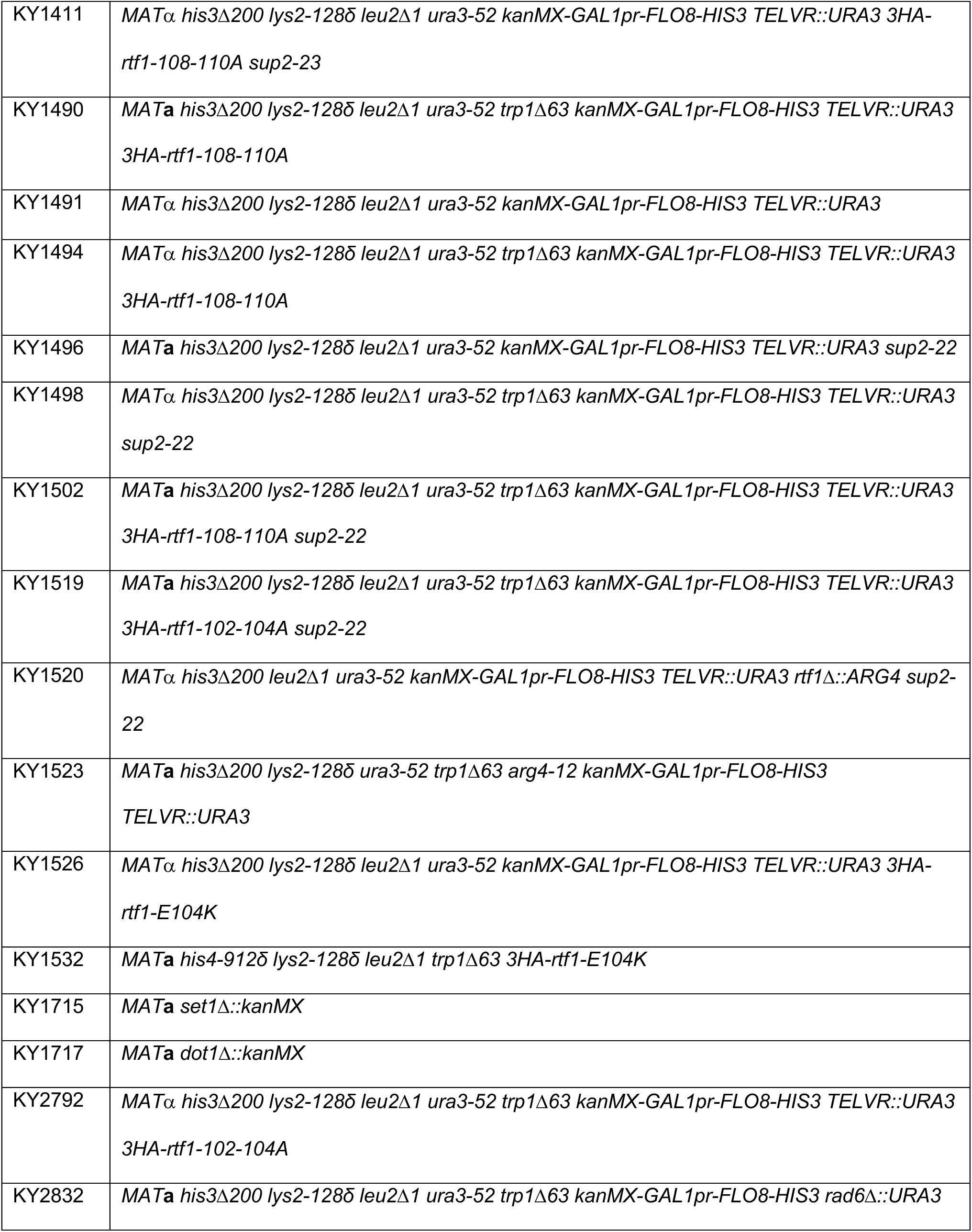
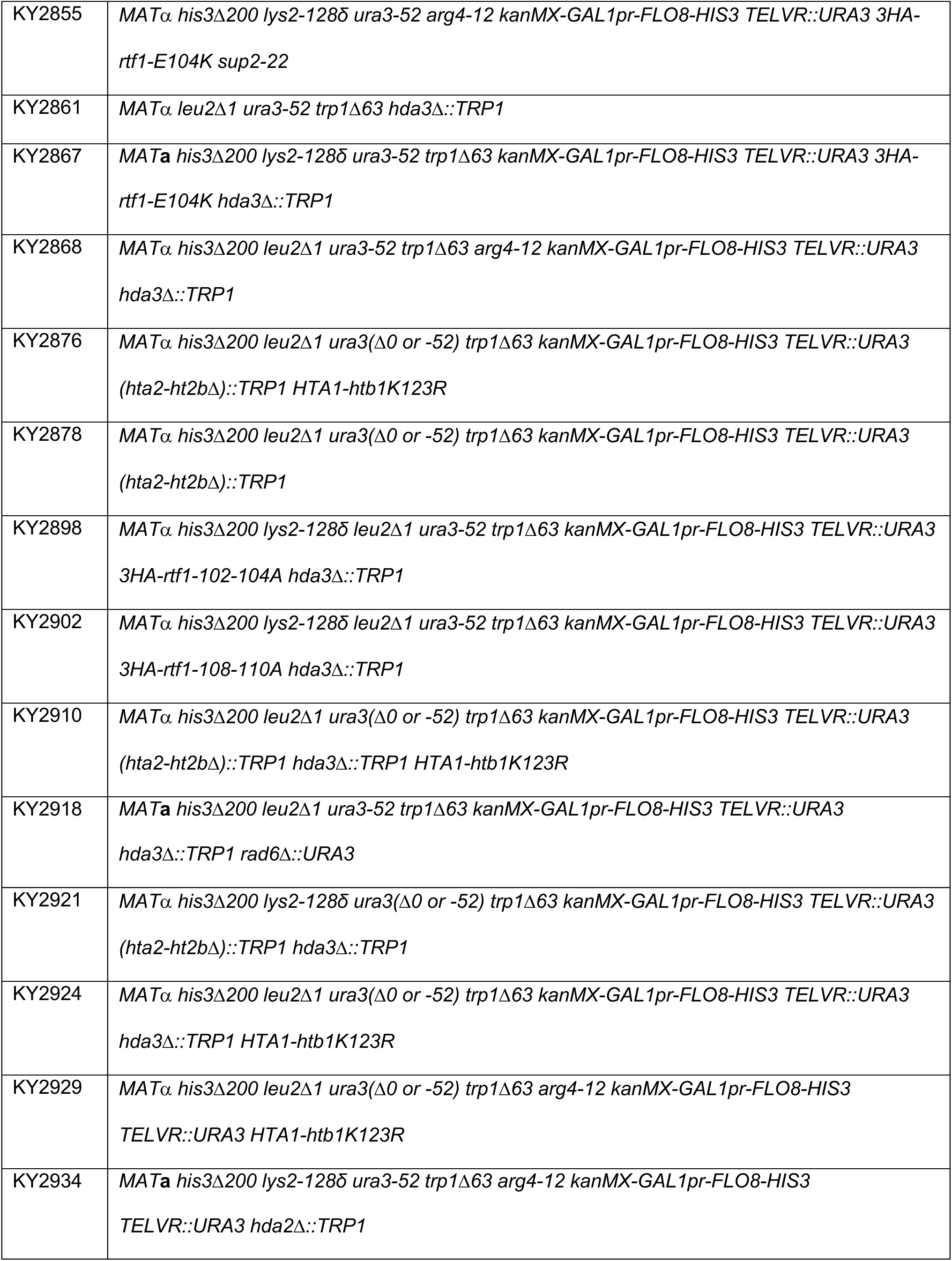
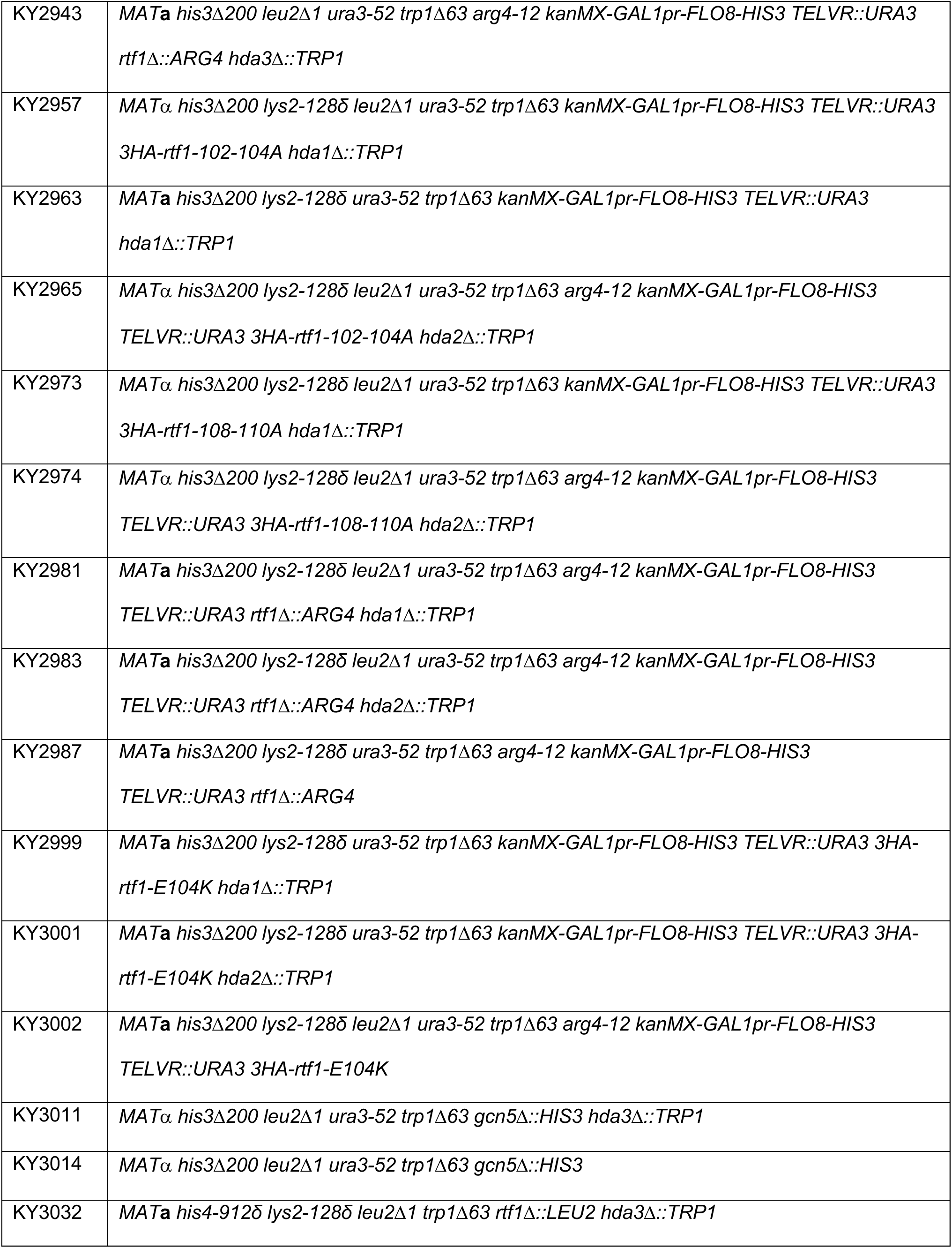
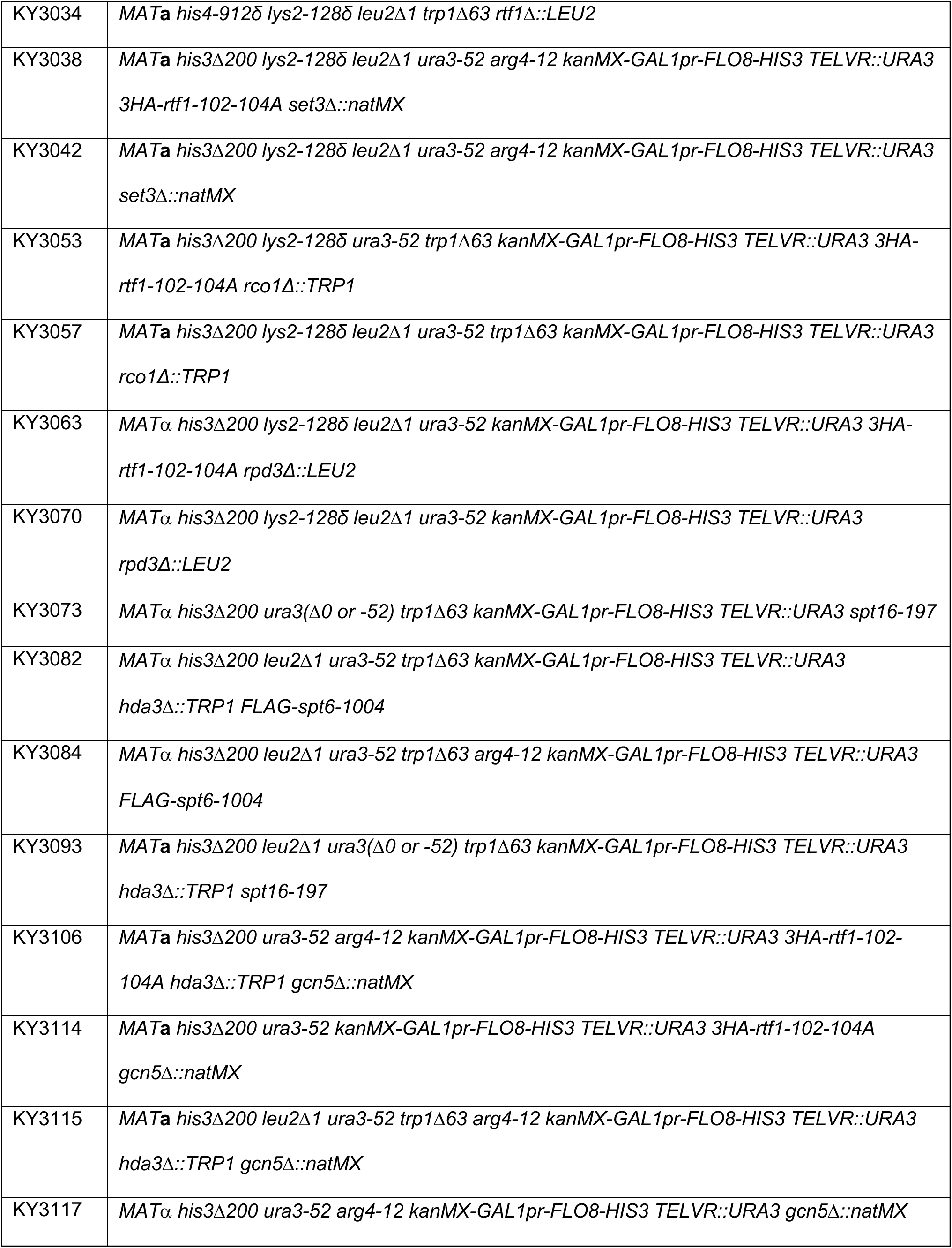
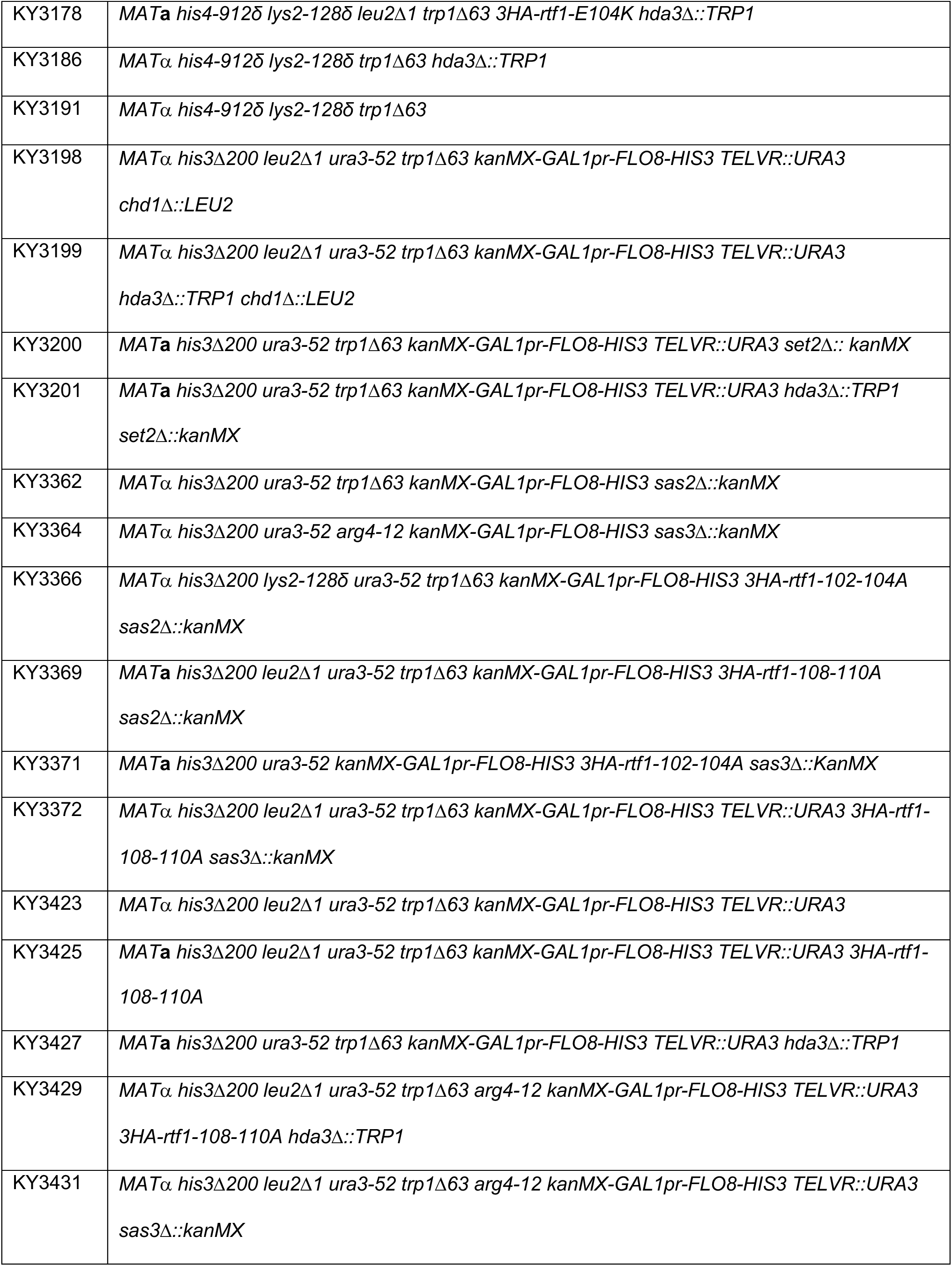
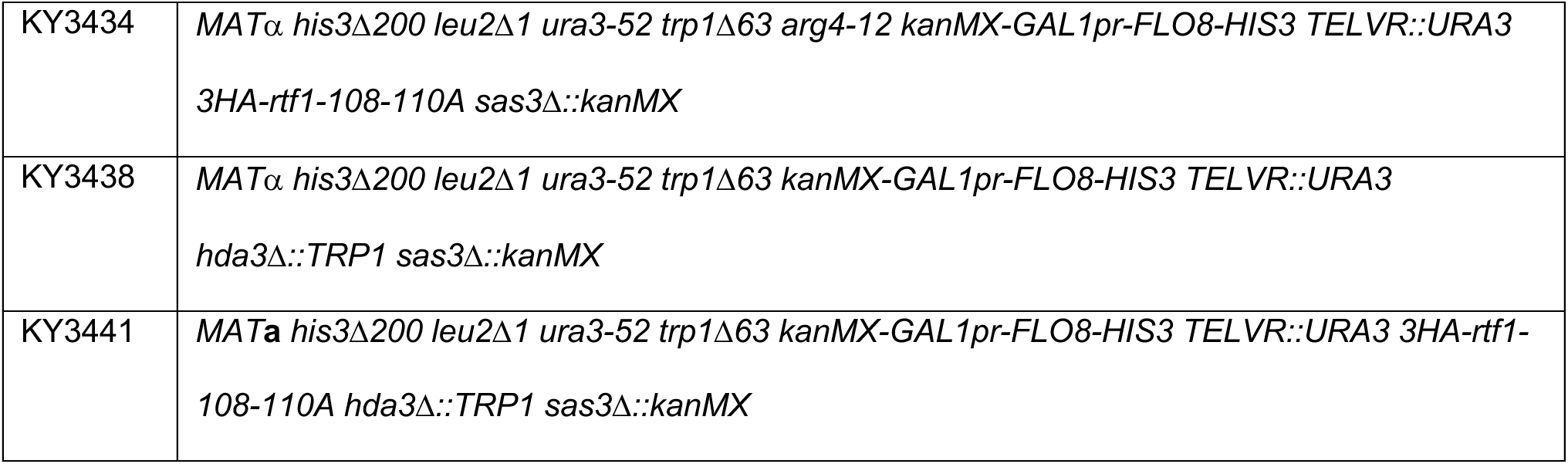
Yeast strains

